# Exposure to early life stress impairs weight loss maintenance success in mice

**DOI:** 10.1101/2023.07.19.549724

**Authors:** Rebecca M Foright, Tara E McQuillan, Jenna M Frick, Paige M Minchella, Brittni M Levasseur, Omar Tinoco, Lauryn Birmingham, Anneka E Blankenship, John P Thyfault, Julie A Christianson

**Affiliations:** Department of Cell Biology & Physiology, University of Kansas Medical Center, Kansas City, Kansas; Kansas Center for Metabolism and Obesity Research; Research Service, Kansas City Veterans Affairs Medical Center, Kansas City, Kansas

**Keywords:** stress, weight regain, exercise, high fat sucrose diet

## Abstract

Early life stress increases obesity risk, but its impact on weight loss maintenance is unknown. Mice underwent neonatal maternal separation (NMS) from 0-3 weeks and were weaned onto high fat sucrose diet (HFSD) from 3-20 weeks. Calorie-restricted weight loss on a low fat sucrose diet (LFSD) occurred over 2 weeks to induce a 20% loss in body weight, which was maintained for 6 weeks. After weight loss, half the mice received running wheels (EX) the other half remained sedentary (SED). Mice were then fed *ad libitum* on HFSD or LFSD for 10 weeks and allowed to regain body weight. NMS mice had greater weight regain, total body weight and adiposity compared to naïve mice. During the first week of refeeding, NMS mice had increased food intake and were in a greater positive energy balance than naïve mice, but total energy expenditure was not affected by NMS. Female mice were more susceptible to NMS-induced effects, including increases in adiposity. NMS and naïve females were more susceptible to HFSD-induce weight regain. Exercise was beneficial in the first week of regain in male mice, but long-term only those on LFSD benefited from EX. As expected, HFSD led to greater weight regain than LFSD.

## INTRODUCTION

Early life stress is increasingly accepted as a driver of weight gain and obesity. Exposure to a range of stressors and trauma can have lasting effects on physiology particularly when the insult occurs during critical periods of development [1]. Clinical research on the topic often uses the Adverse Childhood Experiences (ACEs) questionnaire, which collects self-reported, retrospective data on exposure to a range of potential early life stressors largely encompassing parental behavior and circumstances [2]. Studies consistently find that exposure to early life stress increases obesity risk [3-6]. Notably, even mild forms of early life trauma are associated with an increased risk of obesity [7], type II diabetes [8], sedentary behavior [6], and disordered eating [9].

Rates of obesity and overweight continue to increase worldwide [10]. One of the biggest issues facing those with obesity that want to lose weight is the inability to maintain weight loss long-term. Attempts to lose weight are only transiently effective [11] and 80% of individuals that lose weight regain that lost weight within a year [12]. A key cause for the high recidivism rates is the physiological adaptations that occur following weight loss that elevate appetite and suppress energy expenditure, establishing a strong and persistent biological drive to regain lost weight [13-15]. Although it is increasingly accepted that early life stress exposure contributes to obesity development, there is an overall lack of studies investigating a role for early life stress in the biological drive to regain lost weight.

Regular exercise confers undeniable benefits on overall health and wellbeing [16]. In isolation, exercise has limited benefit in driving weight loss because of the large training volumes needed to elicit the caloric deficits necessary to reduce body weight [17]. There is some clinical evidence this may be a sex-specific response, however, with a greater percentage of men obtaining greater weight loss than women during a 16-week exercise intervention [18]. Following calorie-restricted weight loss, exercise is reported to be a valuable component of successful weight loss maintenance [19] and can counter the biological drive to regain lost weight [19-22].

The present study investigated the role of early life stress on weight loss maintenance success with or without voluntary exercise in both males and females using the mouse model of neonatal maternal separation (NMS). Early life stress can affect both feeding and sedentary behaviors suggesting that both sides of the energy balance equation may contribute to early life stress-induced weight gain. We used indirect calorimetry to dissect the contributions of energy expenditure and energy intake during the first week of *ad libitum* feeding after calorie-restricted weight loss and looked at the effects of stress, sex, diet, and voluntary physical activity on weight regain across 10 weeks. We hypothesized that NMS would drive greater weight regain in both male and female mice. Because we previously found reduced voluntary wheel running distance in early life stressed mice [23], we also hypothesized that NMS would minimize the benefit of exercise on weight loss maintenance.

## METHODS

All procedures were approved by the University of Kansas Medical Center Institutional Animal Care and Use Committee (protocol number 22-04-232) and conformed to the National Institutes of Health *Guide for the Care and Use of Laboratory Animals*. See **Figure 1** for a depiction of the overall study design.

**Figure 1:**
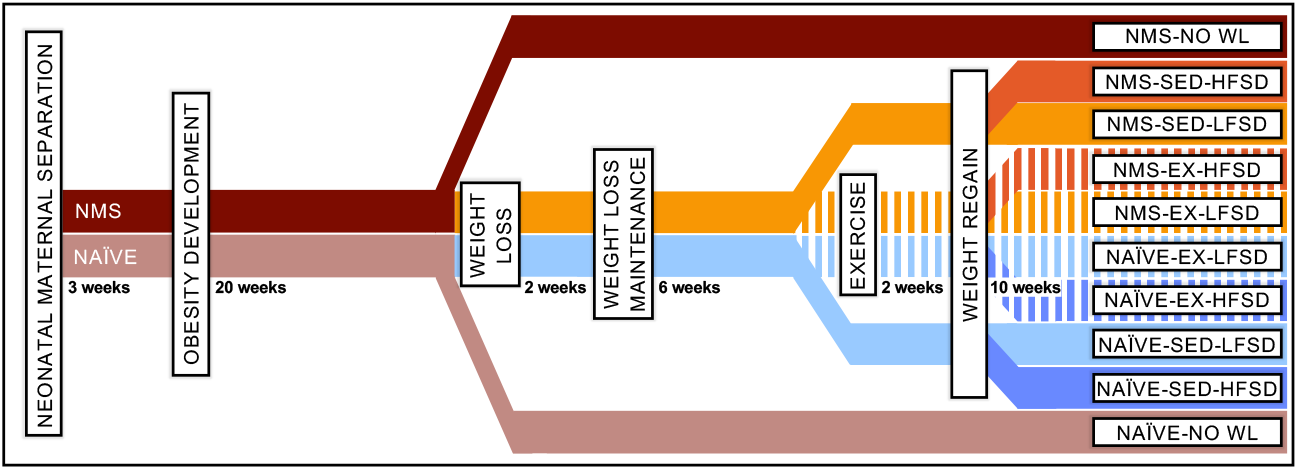
Study Design. Mice underwent neonatal maternal separation from postnatal day 1-21 (NMS) or remained unhandled during this time (naïve). Mice were weaned on to a high fat sucrose diet (HFSD) for 20 weeks. No weight loss (NO WL) control mice stayed on HFSD for the remainder of the study. All other mice underwent calorie-restricted weight loss on a low fat sucrose diet (LFSD) to reduce body weight by 20%. Mice were maintained at the lower body weight for 6 weeks by providing a calorie-limited diet. Half of these mice remained sedentary (SED) all other mice were provided a running wheel in their home cage (EX). After acclimating to the running wheel for 2 weeks, all weight-reduced mice were allowed to feed *ad libitum* on either a LFSD or HFSD. *Ad libitum* feeding continued for 10 weeks. The transition from weight maintained state to *ad libitum* feeding occurred in an indirect calorimetry system.

### Animals

Pregnant C57Bl/6 female mice were ordered from Charles River (Wilmington, MA). Male and female mice used in this study were born and housed in the Laboratory Animal Resource Facility at the University of Kansas Medical Center. All mice had *ad libitum* access to food (except during calorie restriction, see below) and water and were housed on a 12-hour light cycle (0600-1800) at 22ºC.

### Neonatal Maternal Separation (NMS)

Pregnant dams arrived at the animal facility during the last week of gestation. Neonatal maternal separation was performed from postnatal day 1 to 21 (weeks 0-3). Pups were removed from the home cage and placed *en masse* into a clean glass beaker with a thin layer of home cage bedding. The beakers containing the pups were immediately placed in an incubator held at 33ºC and 50% humidity for 180 mins (1100-1400). Dams remained in the home cage during the separation period. Naïve mice remained undisturbed during this time except for the handling necessary for normal husbandry care.

### High Fat Sucrose Diet (HFSD)

All mice were weaned on postnatal day 22, housed with a same sex littermate, and provided high fat sucrose diet (HFSD) (45% kcal fat, 35% kcal carbohydrate (17% sucrose), 20% kcal protein; 4.7 kcal/g; Research Diets, Inc. New Brunswick, NJ; Cat. No. D12451). Fresh HFSD was provided twice weekly.

### Calorie Restricted Weight Loss and Maintenance

After 10 weeks on the HFSD, mice were transitioned to single housing to prepare for the weight loss phase of the study. The no weight loss, control mice were also singly housed at this time to control for the stress induced with social isolation. After 20 total weeks on HFSD, all mice (except no weight loss controls) were switched to low fat sucrose diet (LFSD) (10% kcal fat, 70% kcal carbohydrate (3.5% sucrose), 20% kcal protein; 3.8 kcal/g; Research Diets, Cat. No. D12110704) and calorie restricted. Body weight and food intake were measured daily and adjusted accordingly to produce a 20% decrease in body weight over 2 weeks. Once the target weight loss was achieved, daily body weight and food intake monitoring continued and limited amounts of LFSD were provided to maintain mice at the weight-reduced body weight.

### Voluntary Wheel Running

After 6 weeks of weight loss maintenance, stainless-steel running wheels (STARR Life Sciences Corp., Oakmont, PA) were placed in half of the cages (EX) and remained for the duration of the study. Distance ran was collected during indirect colorimetry housing. Those without running wheels are designated as sedentary (SED) mice.

### Post-Weight Loss *Ad Libitum* Feeding

After 8 weeks of weight loss maintenance, mice were allowed to refeed, *ad libitum*, on either the LFSD or HFSD. Mice remained on their respective diets until study completion 10 weeks later.

### Body Composition

Fat mass and fat-free mass were measured at the end of the study using quantitative magnetic resonance (EchoMRI, Houston, TX).

### Metabolic Monitoring

Mice were housed in an 8-or 16-cage Promethion System (Sable Systems International, Las Vegas, NV). Mice were acclimated to the indirect calorimeter caging for 3 days in the weight-reduced state. After acclimatization, data was collected in the weight reduced state for 3 days and continued for 7 additional days while mice transitioned to the *ad libitum* feeding, weight regain phase. Intake, total energy expenditure (TEE), energy balance, resting energy expenditure (REE), non-resting energy expenditure (NREE), voluntary wheel running distance, and respiratory quotient (RQ) were collected. Weight reduced values were averaged across the 3 collection days. Data from the *ad libitum* refeeding period (7 days) is shown cumulatively across the regain period with the exception of RQ in which a daily average is provided.

### Bomb Calorimetry

Feces were collected during the final two weeks of the study. Fecal samples were dried, pulverized, and pelleted. Fecal pellets were combusted in a 6100 Compensated Jacket Calorimeter (Parr Instrument Company, Moline, IL), filled with oxygen to 40 atm. Ignition was achieved via a high-resistance electrical wire. The calorimeter, upon recording the temperature rise of the surrounding water volume, directly provided the heat of combustion in kcal/g.

### Tissue Collection

At time of euthanasia, mice were overdosed with inhaled isoflurane (>5%). Liver, gastrocnemius, and adipose depots (subcutaneous, mesenteric, retroperitoneal, perigonadal) were dissected and weighed. Visceral adipose was calculated by summing the mesenteric, retroperitoneal, and perigonadal depot weights. While all visceral adipose tissue was collected, only a predefined region of subcutaneous adipose tissue was collected. All tissue specific values are presented as a percentage of total body weight to minimize differences between groups driven solely by differences in total body weight. In doing so, these tissue weights best represent distribution across different tissues and depots.

### Statistics

A 2-way (NMS and sex), 3-way (NMS, sex, exercise or NMS, exercise, diet), or 4-way ANOVA (NMS, sex, exercise, diet) with or without repeated measures was used to test for statistical differences. ANCOVA was run for TEE and REE to determine differences between groups independent of differences in body weight. LSD post hoc analyses were run to determine difference between individual groups. No weight loss mice matched for sex and stress exposure were compared to weight loss-SED-HFSD mice using an independent samples T-test. Significance was set at p<0.05.

## RESULTS

### NMS, female sex, sedentary conditions and HFSD increase weight regain

This study was designed to investigate the impact of NMS, sex, exercise, and diet composition on weight regain following calorie-restricted weight loss (**Figure 1**). Body weight over the course of the entire experiment is shown for male mice (**Figure 2A**) and female mice (**Figure 2B**). NMS mice regained more weight than naïve mice (**Figure 2C**). Female mice regained more weight than males (**Figure 2D**). Sedentary mice regained more weight than exercisers (**Figure 2E**). HFSD-fed mice regained more weight than LFSD-fed mice (**Figure 2F**). Interestingly, female mice regained more weight than male mice on the HFSD while male and female mice regained similar amounts of weight on the LFSD (**Figure 2G**). Exercise was only effective at reducing regain when the mice were on a LFSD (**Figure 2H**). These data show that NMS increases weight regain, and female mice are more susceptible to HFSD-induced weight regain than male mice.

**Figure 2.**
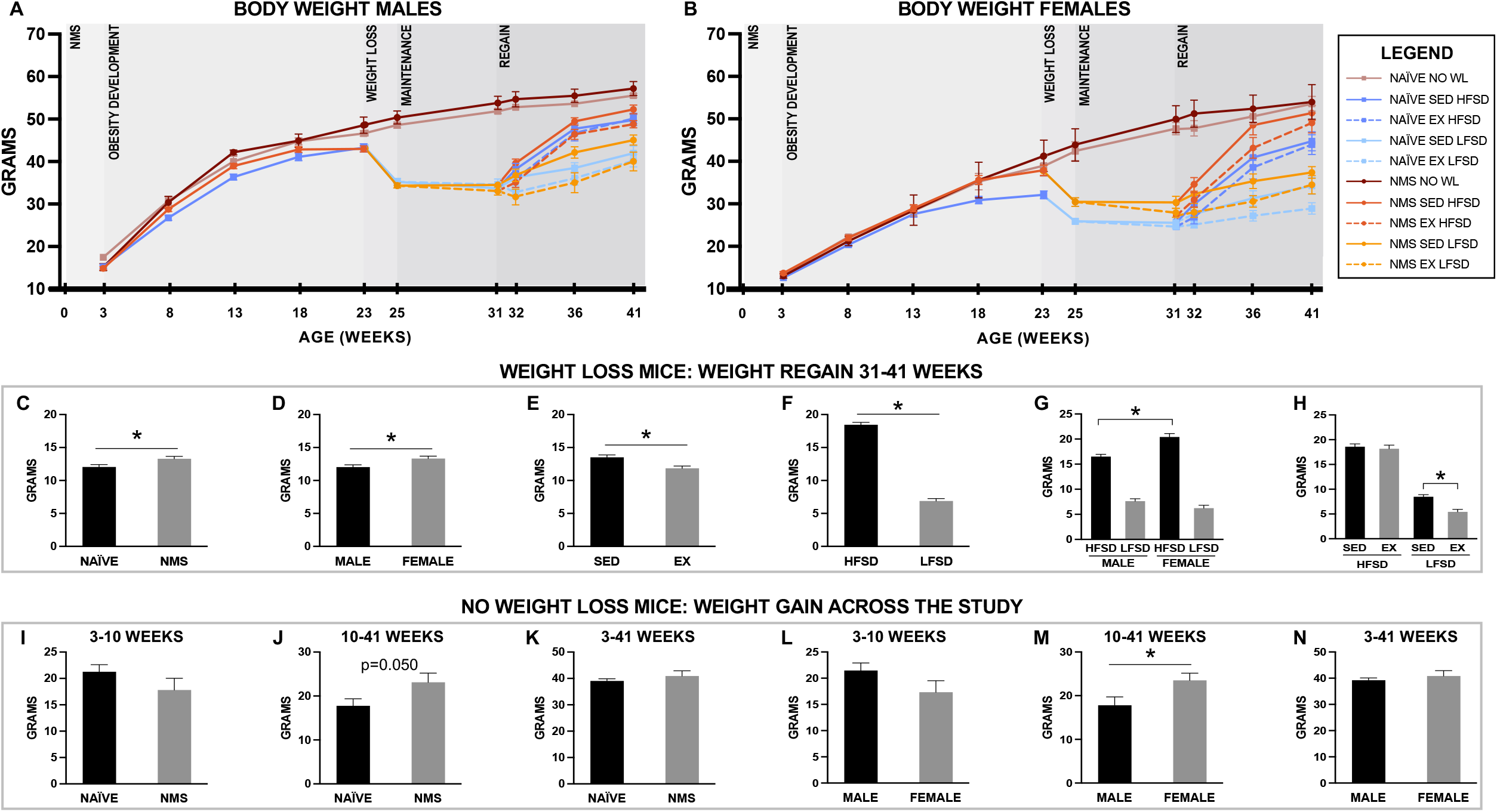
Body weight curves and change in body weight across various timepoints. Body weight across the entire study is plotted for males (A) and females (B). Change in body weight during regain (weeks 31-41) collapsed to show significant differences for stress (C), sex (D), exercise (E), and diet (F) and significant interactions for sex by diet (G) and exercise by diet (H). Change in body weight for no weight loss mice across 3-10 weeks, 10-41 weeks and 3-41 weeks collapsed to show stress (I-K) and sex differences (L-N). Final groups n=6-9. Mean ± SEM displayed. 4-way (C-H) and 2-way (I-N) ANOVAs were used to test significance. Significance was set at p<0.05. Abbreviations: WL, weight loss; HFSD, high fat sucrose diet; LFSD, low fat sucrose diet; NMS, neonatal maternal separation; EX, exercise. SED, sedentary.

For the no weight loss control mice, the effect of stress (**Figures 2I-K**) and sex (**Figures 2L-N**) on HFSD-induced weight gain across the early (3-10 weeks), late (10-41 weeks) and entire (3-41 weeks) study are shown. Across the entire study there were no differences in weight gain between NMS and naïve mice or male and female mice. During the late time point (10-41 weeks), however, it nearly reached statistical significance for NMS mice to gain more weight than naïve mice (**Figure 2J**, p=0.050) and female mice gained significantly more weight than male mice (**Figure 2M**). These data show that while male weight gain slows across long-term HFSD feeding, female weight gain increases (2-way repeated measures ANOVA, time and sex interaction, p<0.050). At the time the study completed, male and female weight gain on HFSD was not different.

### NMS impairs weight loss maintenance success

Weight gain across the 10-week regain period is shown for individual groups for male (**Figure 3A**) and female mice (**Figure 3B**). At the end of the 10-week weight regain period, body weight was significantly increased in NMS, male, SED, and HFSD mice, with significant interaction effects for NMS and sex and sex and diet (**Figure 3C-D**). Body weight at the study completion is shown in **Figures 3C-D** and **Suppl. Table 1**. Female NMS mice (EX-HFSD and SED-LFSD groups) had increased body weight compared to female naïve mice. All naïve female mice, apart from HFSD-SED, weighed less than their male counterparts; however only NMS female mice in the LFSD-EX group weighed significantly less than their male counterparts (**Figure 3D**). EX significantly reduced body weight in male NMS groups, regardless of diet (**Figure 3C**). In contrast, EX only significantly reduced body weight in naïve female mice on LFSD (**Figure 3D**). All LFSD-fed mice weighed significantly less than their NMS/sex/EX-matched counterparts (**Figures 3C-D**).

**Figure 3:**
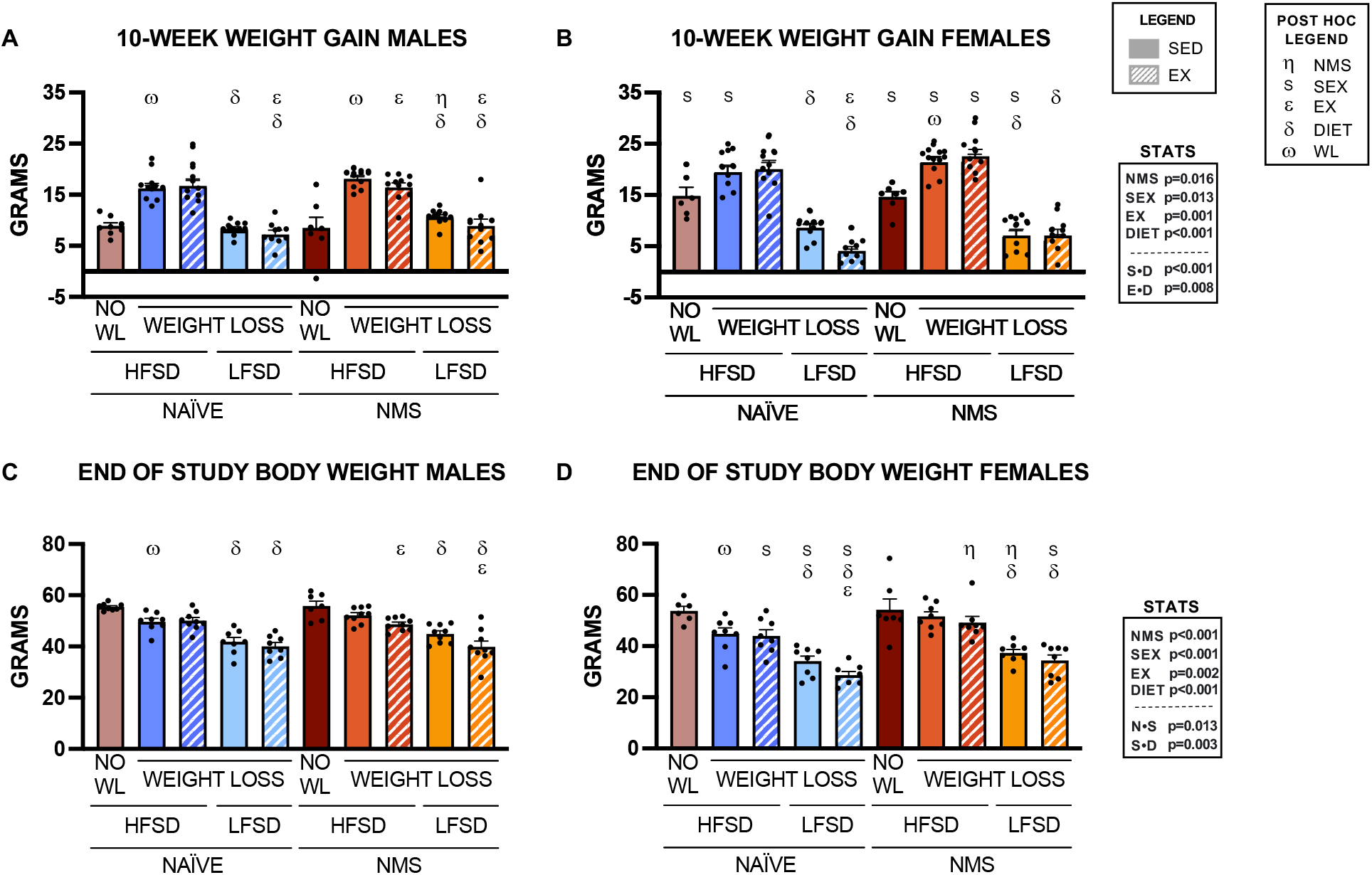
Weight gain during the 10-week regain phase and end of study body weights. Weight gain over the weight regain phase is shown for males (A) and females (B). Ending body weights for males (C) and females (D). Mean ± SEM. N=6-9 per group. 4-way ANOVA was used to test significance (main effects and significant interactions shown in table). Post hoc analysis was determined by LSD (significant comparisons shown in graph). Significance was set at p<0.05. Abbreviations: WL, weight loss; HFSD, high fat sucrose diet; LFSD, low fat sucrose diet; NMS, neonatal maternal separation; EX, exercise; SED, sedentary; S, sex; N, NMS; D, diet.

At the end of the study, male and female naïve-SED-HFSD mice weighed less than their no weight loss control counterparts (**Figure 3C-D**). This demonstrates a lasting benefit of prior weight loss after 10 weeks of *ad libitum* refeeding in naïve mice. Alternatively, male and female NMS-SED-HFSD mice had body weights that were not statistically different from their no weight loss control counterparts (**Figure 3C-D**). These data suggest that prior weight loss had no lasting benefit on body weight in male and female mice that experienced NMS.

### NMS mice have greater total and visceral adiposity

Body composition and tissue-specific weight distribution were measured at the end of the study (**Figure 4, Suppl Table 1**). A significant effect of NMS, sex, EX, and diet were all observed on fat-free mass and adiposity (percent body fat) (**Suppl Table 1**). Significant effect of NMS, EX and diet was observed on fat mass (**Suppl Table 1**). Significant interaction effects of sex and NMS and sex and diet were observed on fat mass and adiposity, with an additional EX and diet interaction effect on adiposity. After 10 weeks of regain, NMS mice not only weighed more, but also had greater adiposity. Female mice on HFSD had significantly greater adiposity than their male counterparts, with the exception of naïve-EX, which was driven by significantly lower fat-free mass (all groups) and by greater fat mass in NMS groups. In addition to significantly reducing weight regain over the 10-weeks, EX reduced adiposity in many groups. Expectedly, all HFSD mice had greater fat mass and adiposity as compared to respective LFSD mice.

**Figure 4:**
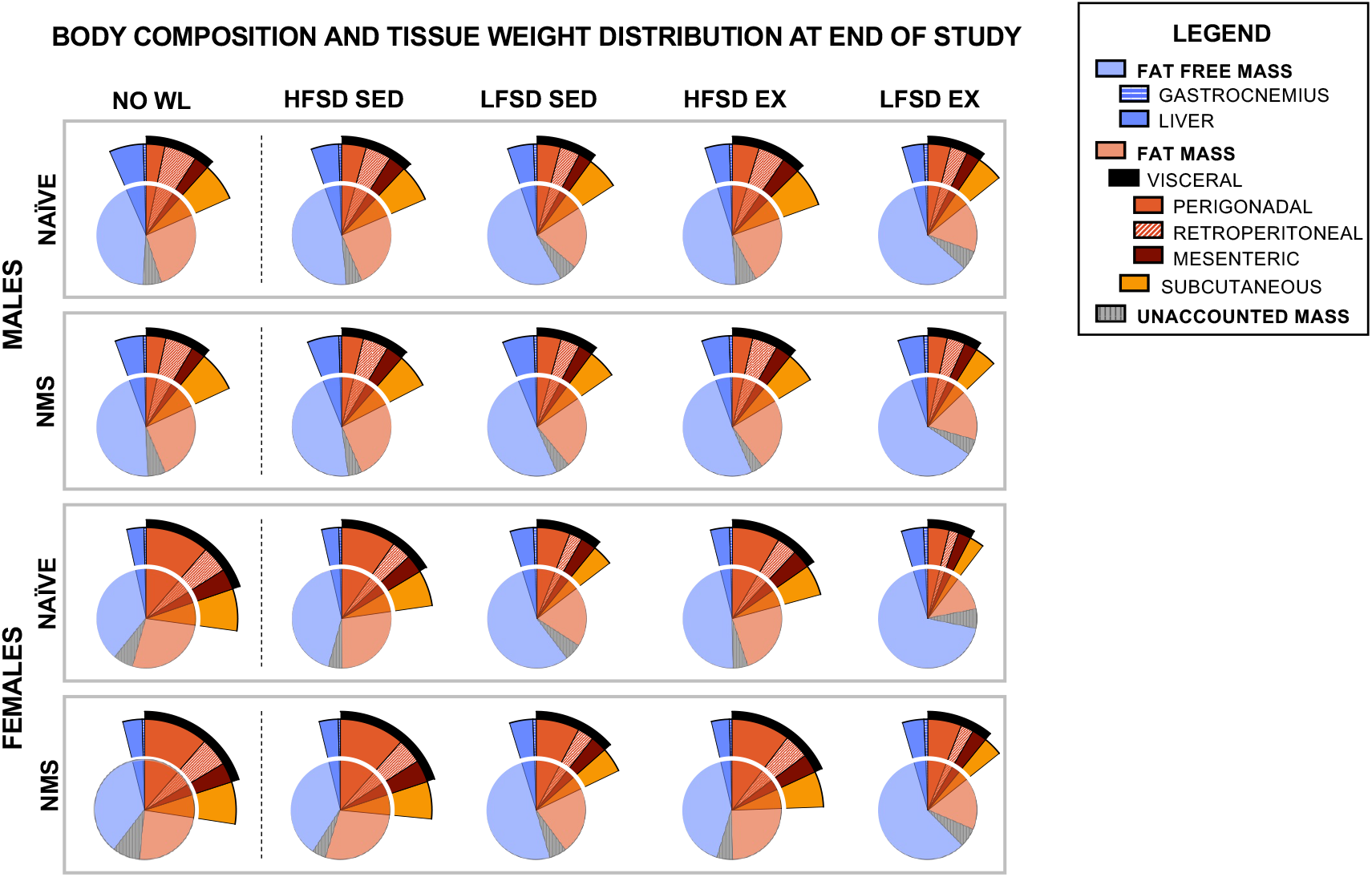
Body composition and tissue weight distribution at the end of the study. Body composition including fat-free mass, fat mass and unaccounted mass (small full circle). Visceral adipose as a percent of total body weight (black pie slice). Tissue-specific weight as a percent of total body weight including gastrocnemius, liver, perigonadal, retroperitoneal, mesenteric and subcutaneous adipose depots (various colors) for mice at the end of the study. For numeric values and statistical significance see Supplemental Table 1. Mean values were used. N=6-9 per group. Abbreviations: WL, weight loss; HFSD, high fat sucrose diet; LFSD, low fat sucrose diet; NMS, neonatal maternal separation; EX, exercise; SED, sedentary.

Tissue-specific weight distribution was affected by all the study variables (**Figure 4, Suppl Table 1**). NMS mice preferentially deposited more weight in perigonadal and retroperitoneal depots resulting in greater visceral adipose tissue weight, although these effects were largely driven by the NMS female mice. Female mice preferentially stored more weight in perigonadal and mesenteric adipose depots while male mice stored relatively more weight in the retroperitoneal depot. Overall, female mice had greater total visceral adipose tissue weight compared to male mice. No weight loss control female mice (naïve and NMS) and NMS-SED-HFSD female mice owed nearly 20% of their total body weight to visceral adipose tissue. EX selectively decreased perigonadal, mesenteric, subcutaneous, and total visceral adipose depot weights but not the retroperitoneal depot weight (**Figure 4, Suppl Table 1**). Expectedly, HFSD increased all adipose depot weights. Post hoc analysis further showed that the retroperitoneal depot weight increased in all groups on HFSD, while the perigonadal depot weight increased in female but not male mice on a HFSD compared to LFSD. Male-NMS mice had heavier livers than naïve male mice and, overall, male mice stored more weight in their livers than female mice especially when on HFSD (**Figure 4, Suppl Table 1**). EX increased relative gastrocnemius weight in several of the exercising groups. Taken together, adiposity and adipose tissue weight distribution occurred in a stress-, sex-, EX-, and diet-specific manner.

### Fecal energy loss is greater on HFSD

Naïve female mice had greater fecal energy loss than naïve male mice which was not seen in NMS mice (**Suppl. Table 1**). Exercising mice on a LFSD had lower fecal energy loss which was not seen in HFSD-EX mice. HFSD-fed mice had greater energy loss than LFSD-fed mice. These differences in fecal energy loss demonstrate that stress, sex, EX, and diet may influence energy absorption beyond changes in energy intake.

### NMS mice had greater food intake and positive energy balance during the first week of regain

While being weight maintained in the weight reduced state, mice were placed in indirect calorimeters to capture the state and transition to *ad libitum* feeding (**Figure 5, Suppl. Table 2**). During the first week of regain, EX resulted in reduced weight regain in all groups except for female mice on HFSD (NMS and naïve) which did not differ from their SED counterpart groups (**Figures 5A-B**). There was a main effect of diet in which all groups gained more weight on the HFSD than their respective LFSD-fed mice (**Figures 5A-B**).

**Figure 5.**
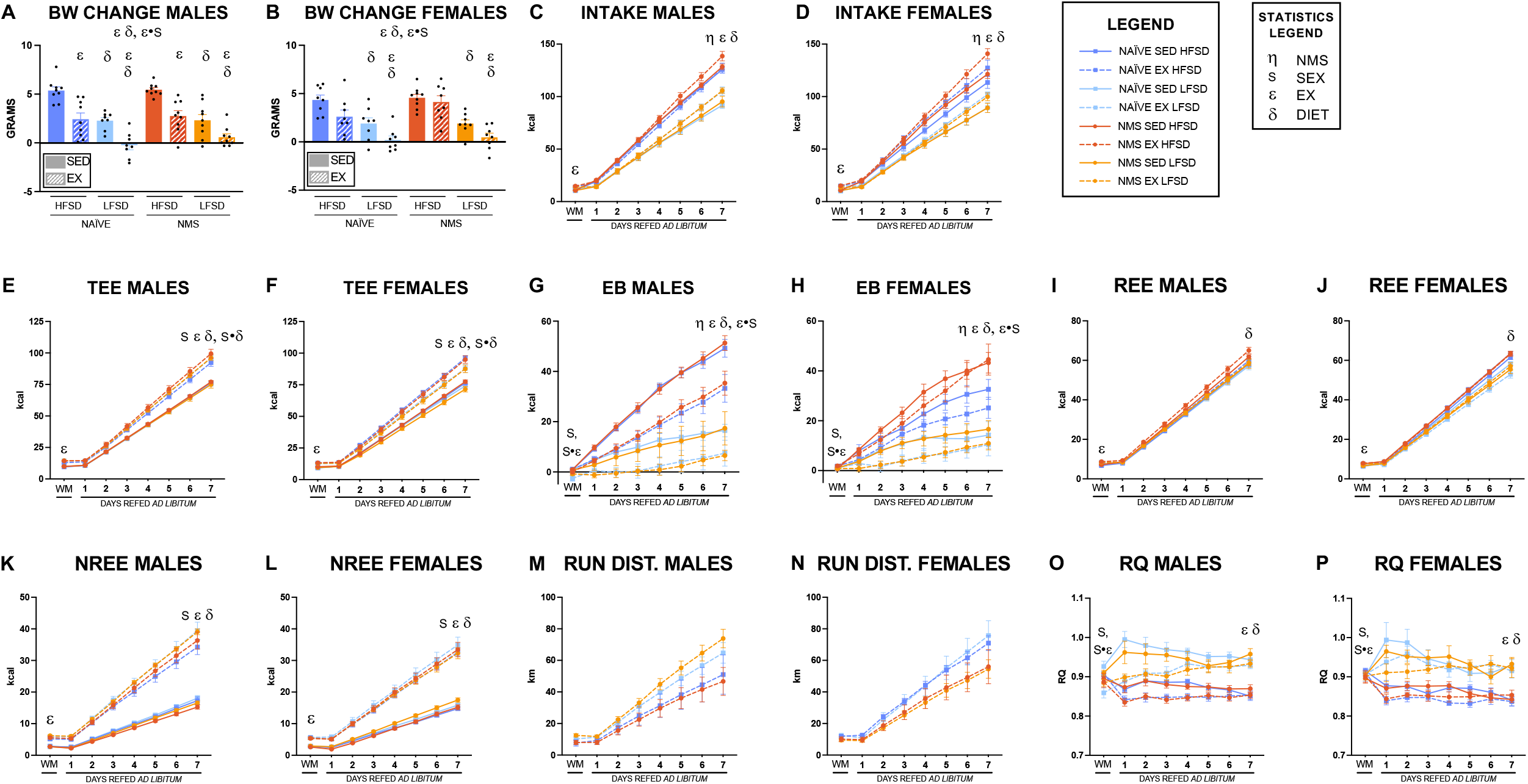
First week of *ad libitum* refeeding. Weight gain over the first 7 days of *ad libitum* refeeding in male (A) and female (B) mice. Energy intake (C & D), total energy expenditure (TEE) (E & F), energy balance (EB) (G & H), resting energy expenditure (REE) (I & J), non-resting energy expenditure (NREE) (K & L), and wheel running distance (M & N) for male and female mice, respectively, in the weight reduced state (average of 3 days) and cumulatively across the first 7 days of *ad libitum* refeeding in male and female mice. Respiratory quotient (RQ) in the weight reduced state and averaged across the first 7 day of *ad libitum* refeeding in male (O) and (P) female mice. Mean ± SEM. N=8-9 per group. 3- and 4-way ANOVAs were run for the weight maintained state (WM) and *ad libitum* day 7, respectively, for energy intake, energy balance, NREE and RQ. For TEE and REE, 3- and 4-way ANCOVAs (body weight) were run at the weight maintained state (WM) and *ad libitum* day 7, respectively. For voluntary wheel running distance, 2- and 3-way ANOVAs were run for the weight maintained state (WM) and *ad libitum* day 7, respectively. Main effect and significant interactions are shown in the graphs. Post hoc analysis was run using LSD and can be found in Supplemental Table 2. Significance was set at p<0.05. Abbreviations: HFSD, high fat sucrose diet; LFSD, low fat sucrose diet; NMS, neonatal maternal separation; EX, exercise; SED, sedentary; TEE, total energy expenditure; EB, energy balance; REE, resting energy expenditure; NREE, non-resting energy expenditure; DIST, distance; RQ, respiratory quotient.

Although NMS mice did not gain significantly more weight than naïve mice during the first week of *ad libitum* refeeding, they consumed more calories and were in a greater positive energy balance (**Figures 5C-D** and **5G-H**). NMS did not affect energy expenditure as total, resting, and non-resting energy expenditure were not altered by NMS exposure (**Figures 5E-F** and **5I-L**).

Spontaneous physical activity also did not differ between NMS and naïve mice (data not shown). In summary, NMS increased food intake and energy balance during the first week of weight regain which, over time, likely contributed to greater total weight regain and increased body weight and adiposity in NMS mice by the end of the study (**Figures 3-4**).

There were interesting differences in energy balance between male and female mice. Male mice were not affected by NMS (p=0.741) but clearly separated by EX (p<0.001) and diet (p<0.001) (3-way ANOVA) (**Figure 5G**). Female mice, however, were not affected by EX (p=0.226) but separated by NMS (p=0.013) and diet (p<0.001) with a significant interaction effect for NMS and diet (p=0.038) such that NMS-HFSD females gained more weight than other groups (3-way ANOVA) (**Figure 5H**).

The first week of weight regain was significantly affected by EX. EX induced greater intake and total energy expenditure (TEE) (**Figure 5E-F**). The increase in TEE with EX was solely attributed to increases in the non-resting energy expenditure component of TEE (**Figure 5K-L**), as EX did not influence resting energy expenditure (**Figure 5I-J**). When running distance was analyzed individually in male mice (**Figure 5M**) there was a significant effect of diet (p=0.036, 2-way ANOVA) such that HFSD-fed male mice ran less than LFSD-fed mice. In female mice (**Figure 5N**), there was a significant effect of NMS (p=0.034, 2-way ANOVA) such that NMS females ran less than naïve females regardless of diet (**Figure 4G**). These sex-specific effects on running distance likely contributed to the sex differences in energy balance and weight regain. Exercise decreased respiratory quotient (RQ) across the regain period (**Figure 5O-P**). Unsurprisingly, diet dramatically affected nearly every variable (**Figures 5C-P**). HFSD increased intake, TEE (females only), energy balance, and REE, and decreased NREE and RQ.

## Discussion

Early life stress increases the risk of developing obesity and metabolic syndrome [3-6, 24]. However, it is unknown how exposure to early life stress may affect weight loss maintenance success. In the present study, we used NMS in mice as a model of early life stress and examined weight regain after calorie-restricted weight loss as compared to naïve mice. We found that NMS increased weight regain across 10 weeks of *ad libitum* feeding and dramatically increased adiposity in regaining NMS females. This work is the first to identify early life stress as a factor contributing to weight regain and showed that female mice were adversely affected to a greater degree than male mice.

Early measurements during the first week of weight regain captured a significant effect of NMS on increased food intake and energy balance but the differences failed to produce increased weight gain over the 7 days. We were not surprised to see that the effects of early life stress on weight regain were more subtle than other well-known factors such as diet and exercise and therefore required additional time for the effects to accumulate over several weeks before weight gain was significantly increased by NMS exposure. We believe this is in large part why early life stress exposure has only recently been accepted as a driver of obesity [24] and has been absent in weight loss maintenance research. We found that these NMS effects albeit more subtle were persistent across the weight regain period and at 10 weeks had significantly increased weight regain.

Exercise significantly decreased weight regain early on and improved weight loss maintenance success long-term in mice on the LFSD, as others have found [25]. Exercising mice on HFSD may have been more susceptible to influence by maladaptive compensations to weight loss which lower energy expenditure to conserve energy and drive weight regain [15]. This was evident during the first week of regain in male mice in the present study which ran lower distances than LFSD-fed male mice. Due to the use of voluntary wheel running in the present study, we hypothesize that the lack of long-term beneficial effects of exercise in HFSD-fed mice was driven by waning running activity and increased sedentary time as is often seen in clinical populations [26].

It was somewhat surprising that early life stress affected female mice to a greater degree than male mice. Our prior work has shown adverse metabolic effects in both male and female mice after NMS [23, 27], although sex differences were not directly tested, as was done in this study. The present study necessitated singly-housing the mice at 10 weeks of age, which we had not previously employed. Following that 10-week point, NMS and naïve male body weights converged. We hypothesize that the social isolation of single housing introduced an additional chronic stressor [28, 29], which may have selectively altered body weight regulation of the naïve male mice, particularly given the presence of HFSD [30]. It is unclear why naïve females did not respond similarly and this sex difference warrants further exploration.

There is a misconception in the field that only male rodents are susceptible to diet-induced obesity. This has historically led many researchers to focus their efforts studying only male rodents despite clinical evidence that women have similar rates of obesity and overweight when compared to men [31]. This work demonstrates increased susceptibility in female mice to diet-induced weight gain after 10 weeks on HFSD and increased susceptibility in weight loss-induced weight regain on HFSD. No weight loss control female mice, including both NMS and naïve groups, displayed profound visceral adiposity with visceral adipose tissue accounting for nearly 20% of their total body weight. The physiological changes that occur to drive these increases in visceral adiposity in only female mice during prolonged HFSD exposure should be identified in future studies.

People with obesity face discrimination across multiple domains. As recently highlighted by Dr. Dakkak, medical stigma of obesity is widespread and prevents people with obesity from seeking care [32]. Furthermore, environmental, economic, political, and systemic racism increase early life stress exposures to those from underserved and historically marginalized communities [33, 34]. This sets the stage for even more discrimination as early life stress-induced weight gain ensues and brings weight bias, stigma, and bullying. It is important that these early life stress drivers of obesity and weight regain be disseminated so that researchers and medical professionals can consider the implications of early life stress-induced obesity when developing new therapeutics and weighing current treatment options.

## CONCLUSIONS

This is the first study to show that early life stress negatively affects weight loss maintenance success through subtle but persistent effects across the entire regain period. This work highlights the importance of future work seeking to understand how early life stress exposure may contribute to challenges in maintaining weight loss in clinical populations.

## Notes

Disclosures: The authors declared no conflict of interest.

### Competing Interest Statement

The authors have declared no competing interest.

## REFERENCES

1. Sutton, E.F., et al., Developmental programming: State-of-the-science and future directions-Summary from a Pennington Biomedical symposium. Obesity (Silver Spring), 2016. 24(5): p. 1018–26.

2. Felitti, V.J., et al., Relationship of childhood abuse and household dysfunction to many of the leading causes of death in adults. The Adverse Childhood Experiences (ACE) Study. Am J Prev Med, 1998. 14(4): p. 245–58.

3. Alvarez, J., et al., The relationship between child abuse and adult obesity among california women. Am J Prev Med, 2007. 33(1): p. 28–33.

4. Bentley, T. and C.S. Widom, A 30-year follow-up of the effects of child abuse and neglect on obesity in adulthood. Obesity (Silver Spring), 2009. 17(10): p. 1900–5.

5. Boynton-Jarrett, R., et al., Child and adolescent abuse in relation to obesity in adulthood: the Black Women’s Health Study. Pediatrics, 2012. 130(2): p. 245–53.

6. Hughes, K., et al., The effect of multiple adverse childhood experiences on health: a systematic review and meta-analysis. Lancet Public Health, 2017. 2(8): p. e356–e366.

7. D’Argenio, A., et al., Early trauma and adult obesity: is psychological dysfunction the mediating mechanism? Physiol Behav, 2009. 98(5): p. 543–6.

8. Thomas, C., E. Hypponen, and C. Power, Obesity and type 2 diabetes risk in midadult life: the role of childhood adversity. Pediatrics, 2008. 121(5): p. e1240–9.

9. Carr, C.P., et al., The Role of Early Life Stress in Adult Psychiatric Disorders: A Systematic Review According to Childhood Trauma Subtypes. The Journal of Nervous and Mental Disease, 2013. 201(12): p. 1007–1020.

10. Federation, W.O., World Obesity Atlas 2023. 2023.

11. Turk, M.W., et al., Randomized clinical trials of weight loss maintenance: a review. J Cardiovasc Nurs, 2009. 24(1): p. 58–80.

12. Wing, R.R. and S. Phelan, Long-term weight loss maintenance. Am J Clin Nutr, 2005. 82(1 Suppl): p. 222S–225S.

13. Fothergill, E., et al., Persistent metabolic adaptation 6 years after “The Biggest Loser” competition. Obesity (Silver Spring), 2016.

14. Maclean, P.S., et al., Biology’s response to dieting: the impetus for weight regain. Am J Physiol Regul Integr Comp Physiol, 2011. 301(3): p. R581–600.

15. Foright, R.M., et al., Is regular exercise an effective strategy for weight loss maintenance? Physiol Behav, 2018. 188: p. 86–93.

16. Garber, C.E., et al., Quantity and Quality of Exercise for Developing and Maintaining Cardiorespiratory, Musculoskeletal, and Neuromotor Fitness in Apparently Healthy Adults: Guidance for Prescribing Exercise. Medicine & Science in Sports & Exercise, 2011. 43(7): p. 1334–1359.

17. Swift, D.L., et al., The Effects of Exercise and Physical Activity on Weight Loss and Maintenance. Prog Cardiovasc Dis, 2018. 61(2): p. 206–213.

18. Donnelly, J.E., et al., Effects of a 16-month randomized controlled exercise trial on body weight and composition in young, overweight men and women: the Midwest Exercise Trial. Arch Intern Med, 2003. 163(11): p. 1343–50.

19. Foright, R.M., et al., Is regular exercise an effective strategy for weight loss maintenance? Physiology & Behavior, 2018. 188: p. 86–93.

20. Higgins, J.A., et al., Resistant starch and exercise independently attenuate weight regain on a high fat diet in a rat model of obesity. Nutrition & Metabolism, 2011. 8(1): p. 49.

21. Levin, B.E. and A.A. Dunn-Meynell, Chronic exercise lowers the defended body weight gain and adiposity in diet-induced obese rats. American Journal of Physiology-Regulatory, Integrative and Comparative Physiology, 2004. 286(4): p. R771–R778.

22. MacLean, P.S., et al., Regular exercise attenuates the metabolic drive to regain weight after long-term weight loss. American Journal of Physiology-Regulatory, Integrative and Comparative Physiology, 2009. 297(3): p. R793–R802.

23. Eller, O.C., et al., Early life stress reduces voluntary exercise and its prevention of diet-induced obesity and metabolic dysfunction in mice. Physiol Behav, 2020. 223: p. 113000.

24. Miller, A.L. and J.C. Lumeng, Pathways of Association from Stress to Obesity in Early Childhood. Obesity (Silver Spring), 2018. 26(7): p. 1117–1124.

25. MacLean, P.S., et al., Regular exercise attenuates the metabolic drive to regain weight after long-term weight loss. Am J Physiol Regul Integr Comp Physiol, 2009. 297(3): p. R793–802.

26. Trost, S.G., et al., Correlates of adults’ participation in physical activity: review and update. Medicine & Science in Sports & Exercise, 2002. 34(12): p. 1996–2001.

27. Frick, J.M., et al., High-fat/high-sucrose diet worsens metabolic outcomes and widespread hypersensitivity following early-life stress exposure in female mice. Am J Physiol Regul Integr Comp Physiol, 2023. 324(3): p. R353–R367.

28. Arakawa, H., Ethological approach to social isolation effects in behavioral studies of laboratory rodents. Behav Brain Res, 2018. 341: p. 98–108.

29. Manouze, H., et al., Effects of Single Cage Housing on Stress, Cognitive, and Seizure Parameters in the Rat and Mouse Pilocarpine Models of Epilepsy. eNeuro, 2019. 6(4).

30. Patterson, Z.R. and A. Abizaid, Stress induced obesity: lessons from rodent models of stress. Front Neurosci, 2013. 7: p. 130.

31. Fryar, C.D.C., Margaret D.; Afful Joseph, revalence of Overweight, Obesity, and Severe Obesity Among Adults Aged 20 and Over: United States, 1960–1962 Through 2017–2018. 2020: NCHS Health E-Stats.

32. Dakkak, M., Fat Shame-Inside and Out. JAMA, 2023.

33. Ash, M. and J.K. Boyce, Racial disparities in pollution exposure and employment at US industrial facilities. Proc Natl Acad Sci U S A, 2018. 115(42): p. 10636–10641.

34. Sacks, V. and D. Murphey, The prevalence of adverse childhood experiences, nationally, by state, and by race/ethnicity. 2018.

