## Supplementary material for "Exposure to early life stress impairs weight loss maintenance success in mice": Suppl. Tables 1-2

| | MALE | | | | | | | | | | FEMALE | | | | | | | | | | SIGNIFICANCE | |
|---|---|---|---|---|---|---|---|---|---|---|---|---|---|---|---|---|---|---|---|---|---|---|
| | NAÏVE | | | | | NMS | | | | | NAÏVE | | | | | NMS | | | | | MAIN EFFECT | INTERACTION |
| | NO WL | PRIOR-WL | | | | NO WL | PRIOR-WL | | | | NO WL | PRIOR-WL | | | | NO WL | PRIOR-WL | | | | | |
| | HFSD | HFSD | | LFSD | | HFSD | HFSD | | LFSD | | HFSD | HFSD | | LFSD | | HFSD | HFSD | | LFSD | | | |
| | SED | SED | EX | SED | EX | SED | SED | EX | SED | EX | SED | SED | EX | SED | EX | SED | SED | EX | SED | EX | | |
| **BW (g)** | 55.5±0.5 | 49.7±1.3* | 50.0±1.2 | 41.8±1.7 | 40.3±1.5 | 55.8±1.9 | 52.2±1.0 | 49.1±1.0 | 45.4±1.2 | 39.3±1.9 | 53.8±1.8 | 44.8±2.3* | 44.0±2.4 | 34.1±2.0 | 28.7±1.4 | 54.3±4.1 | 51.6±1.8 | 49.2±2.4 | 37.3±1.4 | 34.4±2.2 | | |
| NMS | --- | --- | --- | --- | --- | p=0.850 | p=0.153 | p=0.583 | p=0.159 | p=0.697 | --- | --- | --- | --- | --- | p=0.921 | **p=0.036** | p=0.146 | p=0.209 | **p=0.045** | **p<0.001** | NMS*SEX p=0.013<br>DIET*SEX p=0.003 |
| SEX | --- | --- | --- | --- | --- | --- | --- | --- | --- | --- | p=0.324 | p=0.092 | **p=0.047** | **p=0.011** | **p<0.001** | p=0.740 | p=0.785 | p=0.997 | **p=0.001** | p=0.109 | **p<0.001** | |
| EX | | --- | p=0.828 | --- | p=0.502 | | --- | **p=0.043** | --- | **p=0.028** | | --- | p=0.812 | --- | **p=0.046** | | --- | p=0.439 | --- | p=0.272 | **p=0.002** | |
| DIET | | --- | --- | **p=0.003** | **p<0.001** | | --- | --- | **p<0.001** | **p=0.001** | | --- | --- | **p=0.003** | **p<0.001** | | --- | --- | **p<0.001** | **p<0.001** | **p<0.001** | |
| **FAT-FREE (g)** | 27.3±0.4 | 25.5±0.6* | 25.6±0.8 | 24.1±0.5 | 25.4±0.5 | 28.3±0.9 | 27.2±0.7 | 28.0±0.4 | 25.4±0.5 | 25.5±0.6 | 21.0±0.5 | 20.2±0.5 | 21.8±0.5 | 20.1±0.4 | 20.3±0.3 | 21.2±1.2 | 20.9±0.3 | 22.1±0.5 | 20.2±0.2 | 20.9±0.3 | | |
| NMS | --- | --- | --- | --- | --- | p=0.351 | p=0.076 | **p=0.037** | p=0.076 | p=0.934 | --- | --- | --- | --- | --- | p=0.871 | p=0.223 | p=0.728 | p=0.888 | p=0.228 | **p=0.001** | |
| SEX | --- | --- | --- | --- | --- | --- | --- | --- | --- | --- | **p<0.001** | **p<0.001** | **p=0.002** | **p<0.001** | **p<0.001** | **p=0.001** | **p<0.001** | **p<0.001** | **p<0.001** | **p<0.001** | **p<0.001** | |
| EX | | --- | p=0.910 | --- | p=0.076 | | --- | p=0.460 | --- | p=0.900 | | --- | **p=0.024** | --- | p=0.687 | | --- | p=0.070 | --- | p=0.082 | **p<0.001** | |
| DIET | | --- | --- | p=0.082 | p=0.842 | | --- | --- | **p=0.046** | **p=0.006** | | --- | --- | p=0.893 | **p=0.020** | | --- | --- | p=0.091 | p=0.060 | **p=0.005** | |
| **FAT (g)** | 24.9±0.6 | 21.6±0.8* | 21.1±0.9 | 15.4±1.4 | 12.4±1.0 | 24.3±1.4 | 22.7±0.5 | 19.4±0.4 | 17.9±0.8 | 12.0±1.5 | 29.3±1.5 | 22.7±1.9* | 20.1±2.0 | 12.1±1.8 | 6.6±1.1 | 28.2±2.8 | 28.3±1.5 | 24.5±1.8 | 15.1±1.2 | 11.4±2.0 | | |
| NMS | --- | --- | --- | --- | --- | p=0.709 | p=0.254 | p=0.129 | p=0.185 | p=0.802 | --- | --- | --- | --- | --- | p=0.739 | **p=0.031** | p=0.124 | p=0.191 | p=0.053 | **p=0.001** | NMS*SEX p=0.002<br>DIET*SEX p<0.001 |
| SEX | --- | --- | --- | --- | --- | --- | --- | --- | --- | --- | **p=0.033** | p=0.612 | p=0.666 | p=0.168 | **p=0.002** | p=0.242 | **p=0.001** | **p=0.027** | p=0.105 | p=0.840 | p=0.804 | |
| EX | | --- | p=0.691 | --- | p=0.104 | | --- | **p<0.001** | --- | **p=0.006** | | --- | p=0.371 | --- | **p=0.021** | | --- | p=0.125 | --- | p=0.143 | **p<0.001** | |
| DIET | | --- | --- | **p=0.002** | **p<0.001** | | --- | --- | **p<0.001** | **p<0.001** | | --- | --- | **p=0.001** | **p<0.001** | | --- | --- | **p<0.001** | **p<0.001** | **p<0.001** | |
| **ADIPOSITY (%)** | 44.9±0.8 | 43.4±0.8 | 42.0±1.0 | 36.2±2.1 | 30.5±1.4 | 43.5±1.6 | 43.5±0.5 | 39.5±0.3 | 39.2±0.7 | 29.4±2.9 | 54.3±1.2 | 49.9±2.0 | 44.8±2.4 | 34.2±3.5 | 22.0±2.8 | 51.6±1.5 | 54.7±1.0 | 49.5±1.3 | 39.9±1.9 | 31.5±4.1 | | |
| NMS | --- | --- | --- | --- | --- | p=0.469 | p=0.962 | **p=0.043** | p=0.241 | p=0.737 | --- | --- | --- | --- | --- | p=0.178 | p=0.057 | p=0.113 | p=0.172 | p=0.081 | **p=0.004** | NMS*SEX p<0.001<br>EX*DIET p=0.014<br>DIET*SEX p<0.001 |
| SEX | --- | --- | --- | --- | --- | --- | --- | --- | --- | --- | **p<0.001** | **p=0.015** | p=0.305 | p=0.628 | **p=0.020** | **p=0.003** | **p<0.001** | **p<0.001** | p=0.673 | p=0.687 | **p=0.007** | |
| EX | | --- | p=0.311 | --- | **p=0.042** | | --- | **p<0.001** | --- | **p=0.010** | | --- | p=0.129 | --- | **p=0.017** | | --- | **p=0.009** | --- | p=0.085 | **p<0.001** | |
| DIET | | --- | --- | **p=0.011** | **p<0.001** | | --- | --- | **p<0.001** | **p=0.002** | | --- | --- | **p=0.002** | **p<0.001** | | --- | --- | **p<0.001** | **p=0.001** | **p<0.001** | |
| **PERI (%)** | 3.3±0.3 | 4.4±0.3* | 4.7±0.3 | 4.1±0.2 | 4.1±0.3 | 3.5±0.3 | 3.9±0.2 | 3.6±0.2 | 4.3±0.2 | 3.4±0.3 | 11.3±0.4 | 9.6±0.6* | 8.4±0.8 | 5.9±0.8 | 3.8±0.8 | 11.3±0.7 | 11.5±0.3 | 10.3±0.4 | 7.7±0.6 | 6.0±0.9 | | |
| NMS | --- | --- | --- | --- | --- | p=0.644 | p=0.176 | **p=0.006** | p=0.671 | p=0.183 | --- | --- | --- | --- | --- | p=0.937 | **p=0.022** | p=0.051 | p=0.094 | p=0.091 | **p=0.007** | NMS*SEX p<0.001<br>EX*SEX p=0.010<br>DIET*SEX p<0.001 |
| SEX | --- | --- | --- | --- | --- | --- | --- | --- | --- | --- | **p<0.001** | **p<0.001** | **p=0.001** | p=0.052 | p=0.756 | **p<0.001** | **p<0.001** | **p<0.001** | **p=0.001** | **p=0.014** | **p<0.001** | |
| EX | | --- | p=0.512 | --- | p=0.833 | | --- | p=0.402 | --- | p=0.076 | | --- | p=0.229 | --- | p=0.087 | | --- | p=0.057 | --- | p=0.139 | **p=0.001** | |
| DIET | | --- | --- | p=0.449 | p=0.106 | | --- | --- | p=0.251 | p=0.676 | | --- | --- | **p=0.002** | **p=0.001** | | --- | --- | **p<0.001** | **p=0.001** | **p<0.001** | |
| **RP (%)** | 5.8±0.2 | 4.6±0.4* | 4.7±0.2 | 3.5±0.3 | 3.1±0.2 | 4.90±0.3 | 4.5±0.2 | 4.7±0.2 | 3.4±0.1 | 3.2±0.4 | 4.8±0.5 | 3.5±0.3 | 3.6±0.3 | 2.3±0.4 | 1.7±0.3 | 5.1±0.7 | 4.4±0.3 | 4.4±0.4 | 2.9±0.3 | 2.3±0.4 | | |
| NMS | --- | --- | --- | --- | --- | p=0.054 | p=0.742 | p=0.444 | p=0.753 | p=0.759 | --- | --- | --- | --- | --- | p=0.706 | **p=0.028** | p=0.123 | p=0.210 | p=0.315 | **p=0.038** | NMS*SEX p=0.008 |
| SEX | --- | --- | --- | --- | --- | --- | --- | --- | --- | --- | p=0.139 | **p=0.030** | **p=0.010** | **p=0.021** | **p=0.004** | p=0.831 | p=0.865 | p=0.821 | p=0.136 | p=0.164 | **p<0.001** | |
| EX | | --- | p=0.806 | --- | p=0.220 | | --- | p=0.992 | --- | p=0.694 | | --- | p=0.844 | --- | p=0.262 | | --- | p=911 | --- | p=0.249 | p=0.162 | |
| DIET | | --- | --- | **p=0.042** | **p<0.001** | | --- | --- | **p<0.001** | **p=0.040** | | --- | --- | **p=0.023** | **p=0.001** | | --- | --- | **p=0.002** | **p=0.002** | **p<0.001** | |
| **MES (%)** | 2.6±0.1 | 3.1±0.1* | 3.1±0.2 | 2.3±0.1 | 2.4±0.1 | 2.5±0.2 | 3.0±0.2* | 2.9±0.2 | 2.5±0.1 | 2.3±0.2 | 3.7±0.2 | 3.3±0.2 | 3.3±0.2 | 2.8±0.2 | 2.3±0.1 | 3.5±0.2 | 4.0±0.3 | 3.4±0.2 | 2.9±0.2 | 2.6±0.2 | | |
| NMS | --- | --- | --- | --- | --- | p=0.482 | p=0.812 | p=0.188 | p=0.153 | p=0.571 | --- | --- | --- | --- | --- | p=0.572 | p=0.053 | p=0.876 | p=0.636 | p=0.209 | p=0.180 | NMS*SEX p=0.024 |
| SEX | --- | --- | --- | --- | --- | --- | --- | --- | --- | --- | **p=0.002** | p=0.542 | p=0.497 | p=0.066 | p=0.361 | **p=0.002** | **p=0.011** | p=0.058 | p=0.100 | p=0.286 | **p<0.001** | |
| EX | | --- | p=0.803 | --- | p=0.364 | | --- | p=0.343 | --- | p=0.325 | | --- | p=0.804 | --- | p=0.074 | | --- | p=0.092 | --- | p=0.428 | **p=0.025** | |
| DIET | | --- | --- | **p<0.001** | **p=0.004** | | --- | --- | **p=0.027** | p=0.062 | | --- | --- | p=0.132 | **p<0.001** | | --- | --- | **p=0.004** | **p=0.045** | **p<0.001** | |

| | | | | | | | | | | | | | | | | | | | | | | |
|---|---|---|---|---|---|---|---|---|---|---|---|---|---|---|---|---|---|---|---|---|---|---|
| **SQ (%)** | **6.7±0.3** | **6.6±0.6** | **6.9±0.8** | **5.7±0.5** | **4.7±0.4** | **7.2±0.5** | **6.0±0.1** | **5.6±0.4** | **5.1±0.3** | **4.0±0.4** | **7.4±0.4** | **6.4±0.5** | **5.4±0.4** | **3.7±0.4** | **2.6±0.3** | **7.5±0.5** | **6.6±0.5** | **6.3±0.4** | **4.4±0.4** | **3.4±0.5** | | |
| NMS | --- | --- | --- | --- | --- | p=0.404 | p=0.331 | p=0.091 | p=0.271 | p=0.143 | --- | --- | --- | --- | --- | p=0.958 | p=0.718 | p=0.150 | p=0.215 | p=0.179 | p=0.598 | **NMS*SEX p=0.001 DIET*SEX p=0.005** |
| SEX | --- | --- | --- | --- | --- | --- | --- | --- | --- | --- | p=0.145 | p=0.757 | p=0.108 | **p=0.005** | **p=0.001** | p=0.665 | p=0.206 | p=0.124 | p=0.119 | p=0.443 | **p=0.002** | |
| EX | | --- | p=0.759 | --- | p=0.110 | | --- | p=0.146 | --- | **p=0.018** | | --- | p=0.177 | --- | **p=0.046** | | --- | p=0.658 | --- | p=0.114 | **p=0.001** | |
| DIET | | --- | --- | p=0.271 | **p=0.023** | | --- | --- | **p=0.010** | **p=0.009** | | --- | --- | **p=0.001** | **p<0.001** | | --- | --- | **p=0.002** | **p<0.001** | **p<0.001** | |
| **VISCERAL (%)** | **11.7±0.6** | **12.3±0.9** | **12.6±0.8** | **10.0±0.9** | **9.7±0.9** | **10.9±0.7** | **11.4±0.8** | **11.0±0.8** | **10.2±0.8** | **8.8±0.8** | **19.7±0.7** | **16.4±0.9*** | **15.3±0.9** | **10.9±0.9** | **7.8±0.9** | **20.0±0.7** | **19.9±0.9** | **18.1±0.9** | **13.5±0.9** | **10.9±0.9** | | |
| NMS | --- | --- | --- | --- | --- | p=0.275 | p=0.095 | **p=0.007** | p=0.655 | p=0.489 | --- | --- | --- | --- | --- | p=0.847 | **p=0.015** | p=0.092 | p=0.140 | p=0.132 | **p=0.010** | **NMS*SEX p<0.001 EX*SEX p=0.049 DIET*SEX p<0.001** |
| SEX | --- | --- | --- | --- | --- | --- | --- | --- | --- | --- | **p<0.001** | **p=0.003** | p=0.072 | p=0.519 | p=0.190 | **p<0.001** | **p<0.001** | **p<0.001** | **p=0.004** | p=0.290 | **p<0.001** | |
| EX | | --- | p=0.550 | --- | p=0.541 | | --- | p=0.316 | --- | p=0.178 | | --- | p=0.515 | --- | p=0.099 | | --- | p=0.152 | --- | p=0.180 | **p=0.003** | |
| DIET | | --- | --- | **p=0.002** | **p<0.001** | | --- | --- | **p=0.009** | p=0.054 | | --- | --- | **p=0.005** | **p=0.001** | | --- | --- | **p<0.001** | **p=0.002** | **p<0.001** | |
| **LIVER (%)** | **6.0±0.2** | **4.9±0.2*** | **4.3±0.3** | **4.6±0.4** | **3.8±0.1** | **5.0±0.3** | **5.6±0.2** | **5.0±0.3** | **5.6±0.2** | **4.5±0.3** | **3.0±0.1** | **3.0±0.2** | **3.3±0.1** | **4.1±0.2** | **3.9±0.1** | **3.4±0.2** | **3.4±0.1** | **3.2±0.1** | **4.1±0.1** | **3.8±0.1** | | |
| NMS | --- | --- | --- | --- | --- | **p=0.022** | p=0.060 | p=0.171 | **p=0.043** | p=0.077 | --- | --- | --- | --- | --- | p=0.116 | p=0.128 | p=0.306 | p=0.945 | p=0.425 | **p=0.001** | **NMS*SEX p=0.001 EX*SEX p=0.002 p<0.001** |
| SEX | --- | --- | --- | --- | --- | --- | --- | --- | --- | --- | **p<0.001** | **p<0.001** | **p=0.010** | p=0.308 | p=0.648 | **p=0.001** | **p<0.001** | **p<0.001** | **p<0.001** | p=0.060 | **p<0.001** | |
| EX | | --- | p=0.113 | --- | p=0.096 | | --- | p=0.054 | --- | **p=0.032** | | --- | p=0.108 | --- | p=0.450 | | --- | p=0.323 | --- | **p=0.019** | **p<0.001** | |
| DIET | | --- | --- | p=0.410 | p=0.174 | | --- | --- | p=0.980 | p=0.491 | | --- | --- | **p=0.002** | **p=0.001** | | --- | --- | **p<0.001** | **p=0.003** | **p=0.043** | |
| **GAST (%)** | **0.3±0.0** | **0.3±0.0** | **0.3±0.0** | **0.3±0.0** | **0.4±0.0** | **0.3±0.0** | **0.2±0.0** | **0.3±0.0** | **0.3±0.0** | **0.4±0.0** | **0.2±0.0** | **0.3±0.0** | **0.3±0.0** | **0.3±0.0** | **0.4±0.0** | **0.2±0.0** | **0.2±0.0** | **0.2±0.0** | **0.3±0.0** | **0.3±0.0** | | |
| NMS | --- | --- | --- | --- | --- | p=0.400 | p=0.074 | p=0.077 | p=0.120 | p=0.997 | --- | --- | --- | --- | --- | p=0.492 | **p=0.034** | p=0.323 | p=0.452 | p=0.172 | **p=0.010** | **EX*DIET p=0.038 DIET*SEX p=0.023** |
| SEX | --- | --- | --- | --- | --- | --- | --- | --- | --- | --- | **p=0.001** | p=0.525 | p=0.830 | p=0.800 | p=0.246 | p=0.090 | **p=0.001** | **p=0.001** | p=0.448 | p=0.605 | p=0.162 | |
| EX | | --- | p=0.950 | --- | **p=0.032** | | --- | **p<0.001** | --- | **p=0.010** | | --- | p=0.763 | --- | p=0.056 | | --- | **p=0.013** | --- | p=0.157 | **p<0.001** | |
| DIET | | --- | --- | **p=0.022** | **p<0.001** | | --- | --- | **p=0.007** | **p=0.017** | | --- | --- | p=0.640 | **p<0.001** | | --- | --- | **p<0.001** | **p=0.003** | **p<0.001** | |
| **FECAL (kcal/g)** | **3.3±0.0** | **3.3±0.0** | **3.4±0.0** | **3.3±0.0** | **3.2±0.0** | **3.3±0.0** | **3.4±0.0** | **3.4±0.0** | **3.3±0.0** | **3.2±0.0** | **3.3±0.1** | **3.5±0.0*** | **3.4±0.0** | **3.3±0.0** | **3.3±0.0** | **3.4±0.0** | **3.3±0.0** | **3.5±0.0** | **3.3±0.0** | **3.2±0.0** | | |
| NMS | --- | --- | --- | --- | --- | p=0.888 | p=0.150 | p=0.670 | p=0.136 | p=0.075 | --- | --- | --- | --- | --- | p=0.060 | p=0.141 | p=0.732 | p=0.826 | p=0.211 | p=0.637 | **NMS*SEX p=0.020 EX*DIET p=0.009** |
| SEX | --- | --- | --- | --- | --- | --- | --- | --- | --- | --- | p=0.647 | **p=0.027** | p=0.476 | p=0.607 | p=0.052 | p=0.112 | p=0.826 | p=0.499 | p=0.223 | p=0.304 | p=0.063 | |
| EX | | --- | p=0.374 | --- | p=0.055 | | --- | p=0.461 | --- | **p=0.011** | | --- | p=0.601 | --- | p=0.344 | | --- | p=0.145 | --- | p=0.122 | p=0.781 | |
| DIET | | --- | --- | p=0.637 | **p=0.035** | | --- | --- | p=0.080 | **p=0.022** | | --- | --- | **p=0.010** | **p=0.009** | | --- | --- | p=0.244 | **p<0.001** | **p<0.001** | |

Supplemental Table 1: Total body weight, fat-free mass, fat mass and adiposity at the end of the study. Tissue weight as a percent of total body weight at the end of the study for: perigonadal (peri), retroperitoneal (RP), mesenteric (MES), subcutaneous (SQ), and visceral adipose depots, liver, gastrocnemius (GAST) muscle, and fecal energy loss (kcal/g). Between group differences determined by 4-way ANOVA. Main effects and interactions for SEX, NMS, EX, and DIET are displayed in the columns to the right. Post hoc analysis within group was determined by LSD. Post hoc results are displayed in the main body of the table below group values (mean ± SEM). Differences between NO WL/SED/HFSD group and Prior-WL/SED/HFD group within sex and early life stress exposure were determined by t-test and denoted in table with "*". Significance set at P<0.05. Significant results are bolded and colored in purple. Blank cells indicate that no comparison was made. n=6-9 per group. Abbreviations: OB, obese; NMS, neonatal maternal separation; WL, weight loss; SED, sedentary; EX, exercise; HFSD, high fat sucrose diet; LFSD, low fat sucrose diet; BW, body weight; PERI, perigonadal fat; RP, retroperitoneal fat; MES, mesenteric fat; SQ, subcutaneous fat; GAST, gastrocnemius.

| | | MALE | | | | | | | | FEMALE | | | | | | | |
|---|---|---|---|---|---|---|---|---|---|---|---|---|---|---|---|---|---|
| | | NAÏVE | | | | NMS | | | | NAÏVE | | | | NMS | | | |
| | | HFSD | | LFSD | | HFSD | | LFSD | | HFSD | | LFSD | | HFSD | | LFSD | |
| | | SED | EX | SED | EX | SED | EX | SED | EX | SED | EX | SED | EX | SED | EX | SED | EX |
| **INTAKE** | | | | | | | | | | | | | | | | | |
| WEIGHT REDUCED | NMS | --- | --- | --- | --- | p=0.512 | p=0.490 | p=0.567 | p=0.065 | --- | --- | --- | --- | p=0.991 | p=0.665 | p=0.521 | p=0.273 |
| | SEX | --- | --- | --- | --- | --- | --- | --- | --- | p=0.660 | p=0.356 | p=0.287 | **p=0.022** | p=0.975 | p=0.736 | p=0.933 | p=0.854 |
| | EX | --- | **p=0.002** | --- | p=0.405 | --- | **p=0.001** | --- | **p<0.001** | --- | **p<0.001** | --- | **p=0.001** | --- | **p<0.001** | --- | **p=0.002** |
| 7-DAY WEIGHT REGAIN | NMS | --- | --- | --- | --- | p=0.432 | p=0.103 | p=0.566 | p=0.926 | --- | --- | --- | --- | p=0.272 | p=0.144 | p=0.971 | p=0.959 |
| | SEX | --- | --- | --- | --- | --- | --- | --- | --- | p=0.119 | p=0.995 | p=0.699 | p=0.320 | p=0.185 | p=0.698 | p=0.393 | p=0.238 |
| | EX | --- | p=0.681 | --- | **p=0.018** | --- | p=0.075 | --- | p=0.095 | --- | p=0.164 | --- | p=0.073 | --- | **p=0.007** | --- | p=0.083 |
| | DIET | --- | --- | **p<0.001** | **p=0.003** | --- | --- | **p<0.001** | **p<0.001** | --- | --- | **p=0.006** | **p=0.008** | --- | --- | **p<0.001** | **p<0.001** |
| **TOTAL ENERGY EXPENDITURE** | | | | | | | | | | | | | | | | | |
| WEIGHT REDUCED | NMS | --- | --- | --- | --- | p=0.100 | p=0.096 | p=0.727 | p=0.713 | --- | --- | --- | --- | p=0.322 | p=0.649 | p=0.719 | p=0.816 |
| | SEX | --- | --- | --- | --- | --- | --- | --- | --- | p=0.919 | p=0.845 | p=0.411 | p=0.085 | p=0.349 | p=0.082 | p=0.327 | **p=0.038** |
| | EX | --- | **p<0.001** | --- | **p<0.001** | --- | **p<0.001** | --- | **p<0.001** | --- | **p=0.001** | --- | **p=0.009** | --- | **p<0.001** | --- | **p=0.001** |
| 7-DAY WEIGHT REGAIN | NMS | --- | --- | --- | --- | p=0.191 | **p=0.043** | p=0.934 | p=0.917 | --- | --- | --- | --- | p=0.230 | p=0.865 | p=0.625 | p=0.732 |
| | SEX | --- | --- | --- | --- | --- | --- | --- | --- | p=0.584 | p=0.330 | p=0.558 | p=0.093 | p=0.265 | p=0.247 | p=0.571 | p=0.070 |
| | EX | --- | **p<0.001** | --- | **p<0.001** | --- | **p<0.001** | --- | **p<0.001** | --- | **p<0.001** | --- | **p=0.001** | --- | **p<0.001** | --- | **p<0.001** |
| | DIET | --- | --- | p=0.883 | p=0.422 | --- | --- | p=0.454 | p=0.170 | --- | --- | p=0.266 | p=0.057 | --- | --- | **p=0.034** | p=0.071 |
| **ENERGY BALANCE** | | | | | | | | | | | | | | | | | |
| WEIGHT REDUCED | NMS | --- | --- | --- | --- | p=0.857 | p=0.292 | p=0.692 | p=0.171 | --- | --- | --- | --- | **p=0.030** | p=0.923 | p=0.479 | p=0.595 |
| | SEX | --- | --- | --- | --- | --- | --- | --- | --- | p=0.503 | p=0.205 | p=0.482 | **p=0.022** | p=0.354 | p=0.069 | p=0.682 | p=0.077 |
| | EX | --- | p=0.961 | --- | **p=0.012** | --- | p=0.306 | --- | p=0.115 | --- | p=0.248 | --- | p=0.247 | --- | **p=0.036** | --- | p=0.718 |
| 7-DAY WEIGHT REGAIN | NMS | --- | --- | --- | --- | p=0.660 | p=0.780 | p=0.904 | p=0.871 | --- | --- | --- | --- | p=0.084 | **p=0.024** | p=0.719 | p=0.927 |
| | SEX | --- | --- | --- | --- | --- | --- | --- | --- | **p=0.008** | p=0.264 | p=0.716 | p=0.675 | p=0.136 | p=0.261 | p=0.350 | p=0.920 |
| | EX | --- | **p=0.032** | --- | p=0.147 | --- | **p=0.015** | --- | p=0.122 | --- | p=0.225 | --- | p=0.598 | --- | p=0.868 | --- | p=0.205 |
| | DIET | --- | --- | **p<0.001** | **p=0.004** | --- | --- | **p=0.001** | **p<0.001** | --- | --- | **p=0.018** | **p=0.040** | --- | --- | **p<0.001** | **p<0.001** |
| **RESTING ENERGY EXPENDITURE** | | | | | | | | | | | | | | | | | |
| WEIGHT REDUCED | NMS | --- | --- | --- | --- | p=0.152 | **p=0.046** | p=0.372 | p=0.885 | --- | --- | --- | --- | p=0.335 | p=0.495 | p=0.812 | p=.0630 |
| | SEX | --- | --- | --- | --- | --- | --- | --- | --- | p=0.904 | p=0.706 | p=0.547 | p=0.090 | p=0.138 | **p=0.042** | p=0.202 | p=0.347 |
| | EX | --- | **p=0.016** | --- | **p=0.002** | --- | **p=0.001** | --- | **p=0.020** | --- | p=0.387 | --- | p=0.556 | --- | p=0.237 | --- | p=0.316 |
| 7-DAY WEIGHT REGAIN | NMS | --- | --- | --- | --- | **p=0.034** | p=0.073 | p=0.699 | p=0.831 | --- | --- | --- | --- | p=0.274 | p=0.880 | p=0.267 | p=0.144 |
| | SEX | --- | --- | --- | --- | --- | --- | --- | --- | p=0.067 | p=0.149 | p=0.636 | p=0.059 | p=0.266 | p=0.410 | p=0.379 | p=0.443 |
| | EX | --- | p=0.482 | --- | p=0.652 | --- | p=0.121 | --- | p=0.887 | --- | p=0.273 | --- | **p=0.048** | --- | p=0.883 | --- | p=0.579 |
| | DIET | --- | --- | p=0.673 | p=0.570 | --- | --- | p=0.241 | **p=0.017** | --- | --- | p=0.065 | **p=0.002** | --- | --- | **p=0.007** | **p=0.011** |

| | | | | | | | | | | | | | | | | | |
|---|---|---|---|---|---|---|---|---|---|---|---|---|---|---|---|---|---|
| **NON-RESTING ENERGY EXPENDITURE** | | | | | | | | | | | | | | | | | |
| WEIGHT REDUCED | NMS | --- | --- | --- | --- | p=0.977 | p=0.498 | p=0.162 | p=0.473 | --- | --- | --- | --- | p=0.424 | p=0.733 | p=0.837 | p=0.577 |
| | SEX | --- | --- | --- | --- | --- | --- | --- | --- | p=0.878 | p=0.518 | p=0.795 | p=0.484 | p=0.199 | p=0.686 | p=0.245 | p=0.089 |
| | EX | --- | **p<0.001** | --- | **p<0.001** | --- | **p<0.001** | --- | **p<0.001** | --- | **p<0.001** | --- | **p<0.001** | --- | **p<0.001** | --- | **p<0.001** |
| 7-DAY WEIGHT REGAIN | NMS | --- | --- | --- | --- | p=0.156 | p=0.570 | p=0.174 | p=0.903 | --- | --- | --- | --- | p=0.667 | p=0.758 | p=0.097 | p=0.409 |
| | SEX | --- | --- | --- | --- | --- | --- | --- | --- | p=0.082 | p=0.589 | p=0.052 | p=0.242 | p=0.955 | p=0.430 | p=0.338 | **p=0.006** |
| | EX | --- | **p<0.001** | --- | **p<0.001** | --- | **p<0.001** | --- | **p<0.001** | --- | **p<0.001** | --- | **p<0.001** | --- | **p<0.001** | --- | **p<0.001** |
| | DIET | --- | --- | p=0.608 | p=0.167 | --- | --- | p=0.174 | p=0.346 | --- | --- | p=0.193 | p=0.433 | --- | --- | **p=0.045** | p=0.696 |
| **WHEEL RUNNING DISTANCE** | | | | | | | | | | | | | | | | | |
| WEIGHT REDUCED | NMS | | --- | | --- | | p=0.909 | | p=0.093 | | --- | | --- | | p=0.359 | | p=0.074 |
| | SEX | | --- | | --- | | --- | | --- | | p=0.133 | | p=0.131 | | p=0.313 | | p=0.053 |
| 7-DAY WEIGHT REGAIN | NMS | | --- | | --- | | p=0.866 | | p=0.221 | | --- | | --- | | p=0.248 | | p=0.056 |
| | SEX | | --- | | --- | | --- | | --- | | p=0.210 | | p=0.194 | | p=0.524 | | **p=0.040** |
| | DIET | | --- | | p=0.363 | | --- | | **p=0.026** | | --- | | p=0.485 | | --- | | p=0.878 |
| **RESPIRATORY QUOTIENT** | | | | | | | | | | | | | | | | | |
| WEIGHT REDUCED | NMS | --- | --- | --- | --- | p=0.485 | p=0.216 | p=0.293 | p=0.134 | --- | --- | --- | --- | p=0.215 | p=0.559 | p=0.908 | p=0.752 |
| | SEX | --- | --- | --- | --- | --- | --- | --- | --- | p=0.500 | p=0.631 | p=0.229 | **p=0.013** | p=0.954 | p=0.495 | p=0.628 | p=0.213 |
| | EX | --- | p=0.999 | --- | **p=0.003** | --- | p=0.669 | --- | **p=0.043** | --- | p=0.695 | --- | p=0.827 | --- | p=0.810 | --- | p=0.847 |
| 7-DAY WEIGHT REGAIN | NMS | --- | --- | --- | --- | p=0.210 | p=0.987 | p=0.435 | p=0.735 | --- | --- | --- | --- | p=0.925 | p=0.357 | p=0.906 | p=0.827 |
| | SEX | --- | --- | --- | --- | --- | --- | --- | --- | p=0.695 | p=0.312 | p=0.455 | p=0.518 | p=0.052 | p=0.983 | p=0.273 | p=0.674 |
| | EX | --- | p=0.764 | --- | p=0.314 | --- | p=0.233 | --- | p=0.163 | --- | p=0.850 | --- | p=0.545 | --- | p=0.488 | --- | p=0.731 |
| | DIET | --- | --- | **p<0.001** | **p<0.001** | --- | --- | **p<0.001** | **p<0.001** | --- | --- | **p=0.001** | **p=0.008** | --- | --- | **p=0.001** | **p=0.029** |

Supplemental Table 2: Post hoc analysis for intake, total energy expenditure, energy balance, resting energy expenditure, non-resting energy expenditure, wheel running distance and respiratory quotient in the weight reduced state and cumulatively across 7 days of *ad libitum* feeding (RQ is an average on day 7 not cumulative). For main effects and significant interactions see Figure 5. Post hoc analysis within group was determined by LSD. Significance set, P<0.05. Significant results are bolded and colored in purple. Blank cells indicate that no comparison was made. n=6-9 per group. Abbreviations: NMS, neonatal maternal separation; SED, sedentary; EX, exercise; HFSD, high fat sucrose diet; LFSD, low fat sucrose diet.
